# Distinct regulation of tonic GABAergic inhibition by NMDA receptor subtypes

**DOI:** 10.1101/2021.05.30.446187

**Authors:** Kunwei Wu, David Castellano, Qingjun Tian, Wei Lu

## Abstract

Tonic inhibition mediated by extrasynaptic γ-aminobutyric acid type A receptors (GABA_A_Rs) play important roles in the regulation of various brain functions. However, the regulatory mechanisms for tonic inhibition remain largely unknown. Here we report distinct actions of GluN2A- and GluN2B-containing subtypes of NMDA receptors (NMDARs) on tonic inhibition in hippocampal neurons. Mechanistically, GluN2A- and GluN2B-containing NMDARs play differential roles in α5-GABA_A_R internalization. Additionally, GluN2A-, but not GluN2B-, containing receptors are required for the homeostatic potentiation of tonic inhibition. In an acute seizure model induced by kainic acid, tonic inhibition is decreased during acute seizures, while it is increased 24 h later, and these alterations are dependent on the distinct GluN2-containing NMDARs. Collectively, these data reveal a critical link between NMDARs and extrasynaptic GABA_A_Rs in both physiological and pathological conditions.

## INTRODUCTION

γ-aminobutyric acid (GABA), the major inhibitory neurotransmitter in the adult mammalian brain, exerts its fast inhibitory effects by acting on ubiquitously expressed γ-aminobutyric acid type A receptors (GABA_A_Rs). GABA_A_Rs can be classified as mediating phasic or tonic inhibition (Belelli et al., 2009; Farrant and Nusser, 2005). Phasic inhibition is mediated by GABA_A_Rs localized at synapses and activated by GABA released presynaptically in the synaptic cleft, whereas tonic inhibition is mediated by GABA_A_Rs localized outside the synapses (extrasynaptic and perisynaptic) and activated by the low ambient levels of GABA in the extracellular space. Accumulating evidence has revealed that tonic inhibition is involved in a variety of brain functions, including regulating neuronal excitability, neural circuit function, synaptic plasticity, and neuronal development (Belelli et al., 2009; Brickley and Mody, 2012; Farrant and Nusser, 2005; Holter et al., 2010; Lee and Maguire, 2014). Although the molecular and cellular mechanisms regulating phasic inhibition have been extensively studied (Han et al., 2021; Jacob et al., 2008; Luscher et al., 2011; Vithlani et al., 2011), regulatory mechanisms for tonic inhibition remain less clear.

NMDA receptors (NMDARs) are heteromeric complexes assembled from the GluN1 and GluN2 or GluN3 subunit (Chatterton et al., 2002; McBain and Mayer, 1994). The GluN2 subunit has four isoforms (GluN2A to GluN2D), which are differently distributed across the central nervous system (Monyer et al., 1992). In the adult brain, both GluN2A and GluN2B are the predominant GluN2 subunits in the hippocampus (Monyer et al., 1994). GluN2A- and GluN2B-containing NMDARs display differences in their pharmacological and biophysical properties including kinetics, sensitivity to various ligands, permeability to divalent ions, and interactions with intracellular proteins (Vieira et al., 2020). They also have distinct roles in regulating neuronal development (Gambrill and Barria, 2011; Gonda et al., 2020; Sepulveda et al., 2010) and are implicated in pathological conditions such as epilepsy (Chen et al., 2007) and stroke (Chen et al., 2008).

Recent studies have also revealed that NMDARs play an important role in the regulation of GABAergic synapse development and function (Chiu et al., 2018; Gaiarsa, 2004; Gu and Lu, 2018; Gu et al., 2016; Henneberger et al., 2005; Horn and Nicoll, 2018; Marsden et al., 2007; Petrini et al., 2014; Rajgor et al., 2020). However, much less is known about the role of NMDARs in the regulation of tonic GABAergic inhibition. It has been shown that genetic deletion of GluN1, the obligatory subunit of the NMDAR, leads to an enhancement of tonic inhibition in immature hippocampal neurons (Gu et al., 2016). In addition, pathological activation of NMDARs during stroke decreases expression of extrasynaptic δ-subunit-containing GABA_A_Rs and reduces tonic currents in cortical neurons (Jaenisch et al., 2016). However, a systematic investigation of NMDAR regulation of tonic inhibition is lacking. It is also unknown whether the subunit composition of NMDARs is critical for tonic inhibition regulation. Here we employed genetic and pharmacological approaches to demonstrate the differential roles of GluN2A- and GluN2B-containing subtypes of NMDARs in the regulation of extrasynaptic GABA_A_R trafficking and tonic inhibition.

## RESULTS

### Overexpression of GluN2B inhibits tonic inhibition

To assess the role of GluN2 subunits in modulation of tonic inhibition, we first overexpressed GluN2A or GluN2B in cultured hippocampal neurons at DIV11, and then measured miniature inhibitory postsynaptic currents (mIPSCs) and tonic inhibitory currents at DIV14 (Figure 1A). We observed that tonic currents were significantly reduced in neurons overexpressing GluN2B, whereas overexpression of GluN2A had no effect on tonic currents, compared to control neurons overexpressing GFP (Figure 1B and 1C). However, neither GluN2A nor GluN2B overexpression had a significant effect on mIPSCs (Figure S1), indicating a subunit specific function of GluN2 in regulating tonic, but not phasic, inhibition. Previous work has shown that extrasynaptic α5-GABA_A_Rs mediate the majority of tonic inhibition in hippocampal pyramidal neurons (Caraiscos et al., 2004; Glykys et al., 2008). To investigate whether overexpression of GluN2 subunits regulated the abundance of surface α5-GABA_A_Rs, we performed α5-GABA_A_R immunostaining assays in cultured hippocampal neurons. Consistent with the effect on tonic currents (Figure 1B), overexpression of GluN2B, but not GluN2A or the control, GFP, reduced the surface expression of α5-GABA_A_Rs (Figure 1C).

**Figure 1.**
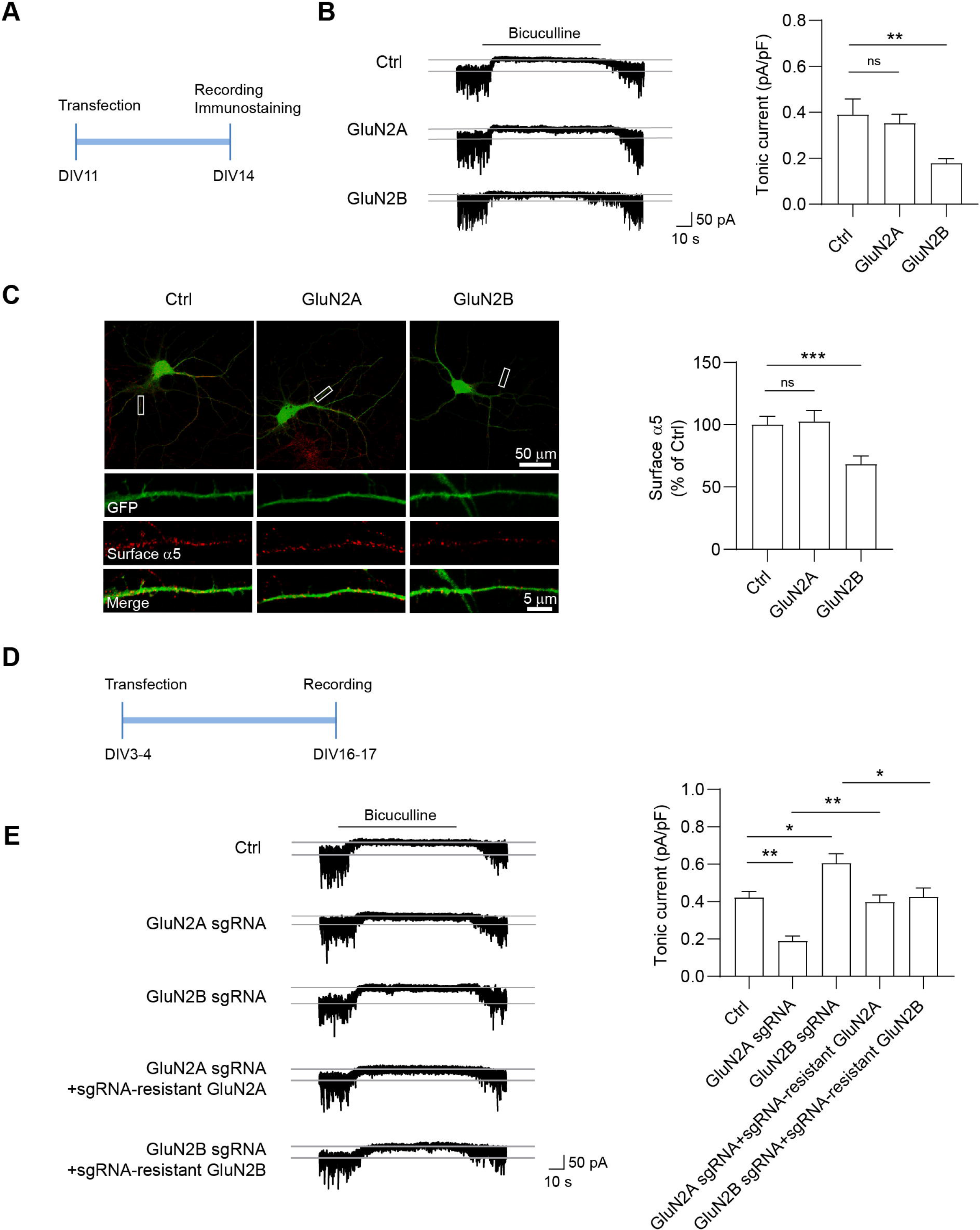
Overexpression of GluN2B inhibits tonic inhibition. (A) Overexpression experiment design. Neurons were transfected at 11 days *in vitro* (DIV11) for 72 h and then recorded for tonic inhibitory currents at DIV14. (B) Representative traces and summary graphs showing that GluN2B, but not GluN2A overexpression decreased tonic currents in cultured hippocampal neurons. (n = 11–13 for each group, one-way ANOVA test with Dunnett’s multiple comparisons test) (C) Immunostaining and summary graphs showing that GluN2B but not GluN2A overexpression decreased surface α5 expression in cultured hippocampal neurons. (n = 28-31 for each group, one-way ANOVA test with Dunnett’s multiple comparisons test). (did not show where the amplified region comes from in the images) (D) Knockout (KO) experiment design. Neurons were transfected at DIV3-4 and then recorded for tonic currents at DIV16-17. (E) Representative traces and summary graphs showing that GluN2A KO decreased tonic currents, whereas GluN2B KO increased tonic currents. The changes in tonic currents induced by either GluN2A or GluN2B KO were restored back to the control level by co-expression of corresponding sgRNA-resistant constructs, respectively. (n = 8–10 for each group, one-way ANOVA test Tukey’s multiple comparisons test) *p < 0.05 and **p < 0.01. All data are presented as mean ± SEM.

### Knockout of GluN2A and GluN2B differentially regulate tonic inhibition

To complement the overexpression experiments, we developed single-guide RNAs (sgRNAs) to perform single-cell genetic deletion of endogenous GluN2A or GluN2B in cultured hippocampal neurons and then measured tonic inhibitory currents (Figure 1D). The knockout (KO) efficacy was confirmed by Western blot in co-transfected HEK293 cells (Figure S2A and S2B) as well as by NMDA mEPSC and NMDA-evoked whole-cell current recordings in cultured neurons transfected with both sgRNAs (Figures S2C, S2D and S2E). We found that whereas sgRNA-mediated KO of GluN2A significantly decreased tonic currents, GluN2B KO increased tonic currents, compared with control neurons expressing empty vector (Figure 1E). Importantly, the changes in tonic currents induced by either GluN2A or GluN2B KO were restored back to the control level by co-expression of corresponding GluN2 sgRNA-resistant constructs (Figure 1E), suggesting that these effects were due to loss of GluN2A or GluN2B, but not off-target effects.

### Pharmacological suppression of GluN2A- and GluN2B-containing receptors modulateα5-GABA_A_R surface expression and tonic inhibition

To corroborate our genetic findings, we next employed pharmacological treatment to examine the role of GluN2A and GluN2B in the regulation of tonic inhibition. Specifically, we treated cultured hippocampal neurons at DIV15 with NVP-AAM077 (NVP, GluN2A-preferring antagonist; 100 nM), ifenprodil (Ifen, GluN2B-preferring antagonist; 5 μM) or APV (broad-spectrum NMDAR antagonist; 100 μM) for 24 h, and then measured tonic inhibitory currents. We found that Ifen increased surface α5 expression and tonic inhibitory currents, whereas NVP induced opposite effects (Figures 2B and 2C). In addition, APV had little effect on surface α5 expression and tonic currents (Figures 2B and 2C), indicating that blockade of both of GluN2A- and GluN2B-containing NMDARs exerts a combinatory effect leading to little change in tonic inhibition. Taken together, these results suggest the differential regulation of α5 surface expression and tonic inhibition by GluN2A- and GluN2B-containing receptors.

**Figure 2.**
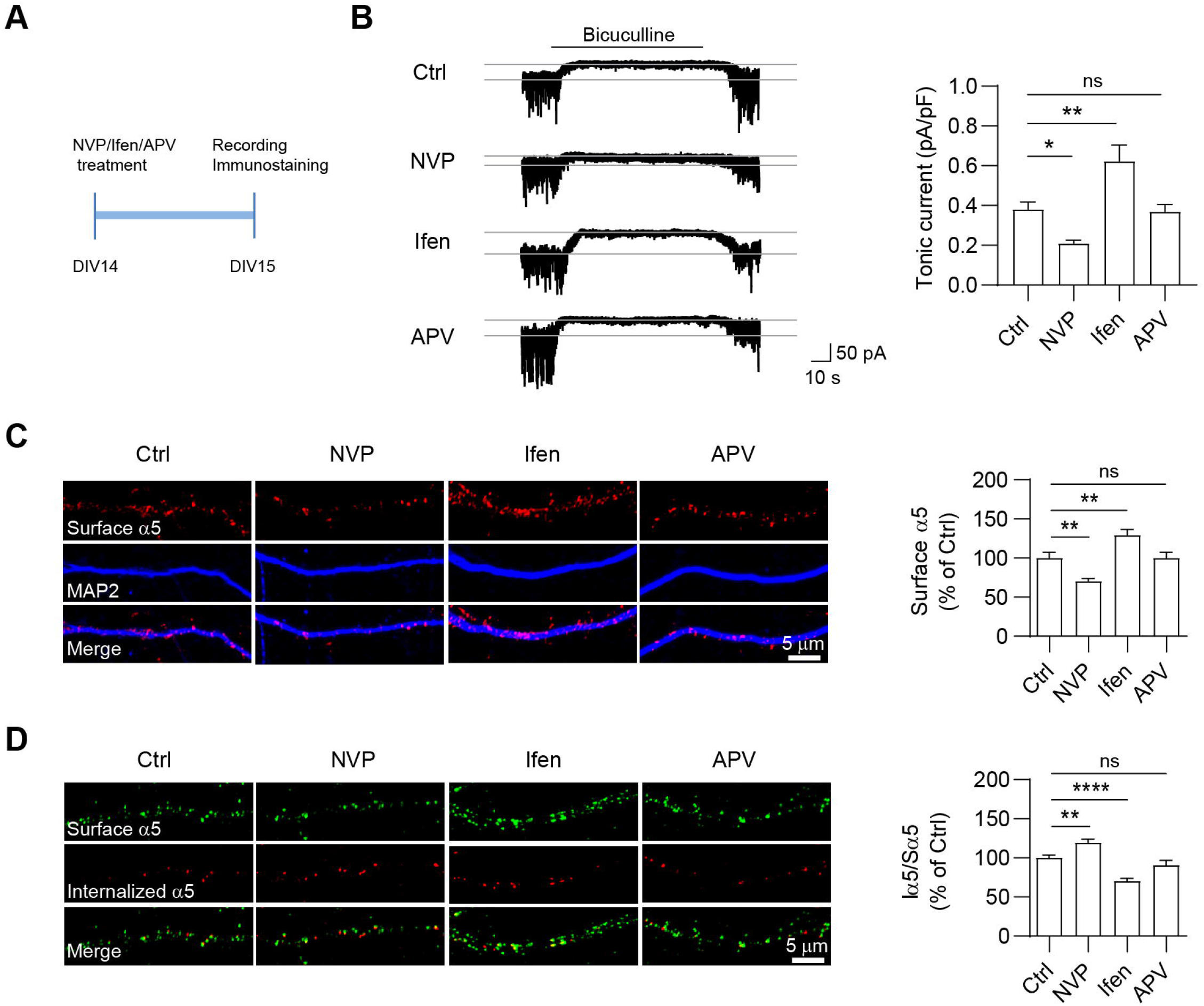
Pharmacological suppression of GluN2A- and GluN2B-containing receptors regulate tonic inhibition and α5-GABA_A_R internalization. (A) Experimental design. Hippocampal neurons at DIV 14 were treated with NVP (GluN2A-preferring antagonist NVP-AAM077, 100 nM), Ifen (GluN2B-preferring antagonist ifenprodil, 5 μM), or APV (broad-spectrum NMDAR antagonist, 100 μM) for 24 h and then recorded for tonic currents at DIV15. (B) Representative traces and summary graphs showing that 24-h Ifen treatment increased tonic currents, whereas 24-h NVP treatment decreased tonic currents in cultured hippocampal neurons. (n = 10-11 for each group, one-way ANOVA with Dunnett’s multiple comparisons test) (C) Immunostaining and summary graphs showing 24-h Ifen treatment increased surface α5 expression, whereas 24-h NVP treatment decreased surface α5 expression in cultured hippocampal neurons. (n = 33-43 for each group, Kruskal-Wallis test with Dunnett’s multiple comparisons test) (D) Endocytosis assay of α5-GABA_A_Rs in hippocampal cultures. Surface α5-GABA_A_Rs (Sα5) were labeled in green, and internalized α5-GABA_A_Rs (Iα5) were in red. Bar graphs in the right showing that 24-h NVP treatment increased α5 internalization, whereas 24-h Ifen treatment decreased α5 internalization. (n = 15-16 for each group, one-way ANOVA with Dunnett’s multiple comparisons test). *p < 0.05, **p < 0.01 and ****p < 0.0001. All data are presented as mean ± SEM.

During development, NMDARs undergo a developmental switch from those containing primarily the GluN2B subunit to primarily the GluN2A subunit in neocortex and hippocampus (Dong et al., 2006; McKay et al., 2018; Sheng et al., 1994). Consistent with these previous studies, we confirmed the developmental switch from primarily GluN2B-containing receptors to primarily GluN2A-containing receptors in cultured hippocampal neurons (Figure S3A). In addition, both GABA_A_R α5 subunit expression and tonic currents were increased during development within the first month (Figures S3A and S3B), suggesting a possible link between developmental switch of NMDARs subunits and developmental increase of α5-GABA_A_R expression within the first month. Thus, we tested the effects of GluN2A and GluN2B antagonists on tonic inhibition in immature and more differentiated neurons. In immature neurons at DIV7-8, Ifen or APV increased surface α5 expression and tonic currents, whereas NVP had little effect (Figures S3C and S3D), consistent with higher expression of GluN2B in developing, immature neurons. By contrast, in more differentiated neurons at DIV25-26, NVP and APV decreased surface α5 expression and tonic currents, whereas Ifen had little effect (Figures S3C and S3D). These results suggest that regulation of tonic inhibition by GluN2A and GluN2B is developmentally dependent.

### Pharmacological suppression of GluN2A- and GluN2B-containing receptors regulateα5-GABA_A_R internalization

Considering that opposite actions of GluN2A- and GluN2B-containing receptor blockade in α5-GABA_A_R surface expression and tonic currents were observed at 2-3 weeks neurons, we thus investigated the mechanism underlying the distinct effects at this developmental stage in the following experiments. The abundance of surface α5-GABA_A_Rs is controlled by the balance of receptor endocytosis and exocytosis. Since surface expression of α5-GABA_A_Rs is differentially regulated by NMDAR GluN2 subunits, we hypothesized that α5 endocytosis and/or exocytosis might be critically regulated by GluN2A- or GluN2B-containing NMDARs. To test this hypothesis, we first performed antibody-feeding experiments to label surface and internalized endogenous α5 in live hippocampal neurons at DIV15 and examined α5 endocytosis. We found that 24-h NVP treatment increased α5 internalization, whereas 24-h Ifen treatment decreased α5 internalization (Figure 2D). In addition, 24-h APV treatment did not alter α5 internalization (Figure 2D). These results suggest that GluN2A and GluN2B have distinct role in regulating α5-GABA_A_R internalization at this developmental stage. Next, we combined fluorescence recovery after photobleaching (FRAP) with fluorescence loss in photobleaching (FLIP) to investigate exocytosis of superecliptic pHluorin-tagged α5 (SEP-α5) to measure the receptor exocytosis. In this experiment, repetitive photobleaching occurred at dendritic regions bilateral to the central FRAP area, thus excluding laterally diffusing SEP-α5 to the central area and allowing the measurement of newly exocytosed SEP-α5 (Figure S3E). We found that neurons under different treatments exhibited similar fluorescence recovery after photobleaching (Figure S3F), indicating that neither GluN2A- nor GluN2B-containing receptors regulate α5 exocytosis.

### GluN2A-containing receptors are required for homeostatic potentiation of tonic inhibition

Chronic pharmacological manipulation of neuronal activity can induce homeostatic adaptions of excitatory and inhibitory transmission, which are powerful mechanisms controlling neuronal excitability and neural network function (Turrigiano, 2012). Our recent work has demonstrated that tonic inhibition in hippocampal neurons exhibited homeostatic plasticity (Wu et al., 2021). Consistent with this study, we found that surface α5 expression and tonic inhibition were increased following 48-h bicuculline treatment (Figure 3). To examine the role of GluN2A- and GluN2B-containing NMDARs in homeostatic plasticity of tonic inhibition, we treated hippocampal neurons at DIV16 with bicuculine (40 μM) and 24 h later NVP (100 nM), Ifen (5 μM) or APV (100 μM) was administered for 24 h before recording. We found that bicuculline-induced effects were significantly diminished by treatment of NVP and APV, but not by Ifen (Figure 3). These observations support a necessary role for GluN2A-, but not GluN2B-, containing NMDARs in homeostatic potentiation of tonic inhibition.

**Figure 3.**
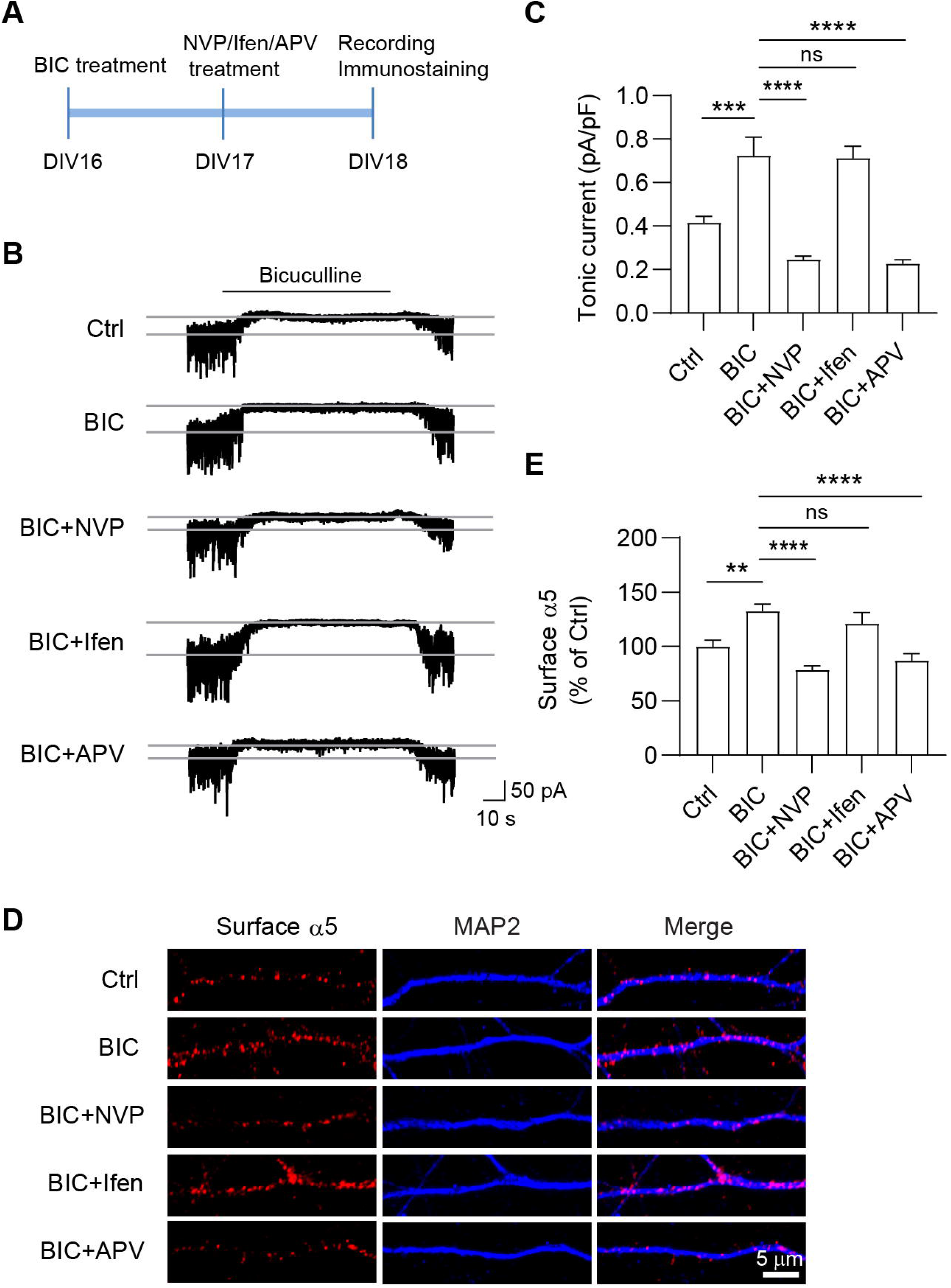
GluN2A-contaning receptors are required for homeostatic potentiation of tonic inhibition. (A) Experimental design. Hippocampal neurons at DIV 16 were treated with bicuculline (BIC, 40μM), and at DIV 17 were treated with NVP (100 nM), Ifen (5 μM), or APV (100 μM) for 24 h before recording. (B-C) Representative traces (B) and summary graphs (C) showing that NVP and APV treatment abolished BIC-induced potentiation of tonic currents in cultured hippocampal neurons. (n = 10 for each group, one-way ANOVA with Tukey’s multiple comparison test). (D-E) Immunostaining (D) and summary graphs (E) showing that NVP and APV treatment abolished BIC-induced potentiation of surface α5 expression in cultured hippocampal neurons. (n = 20-26 for each group, Kruskal-Wallis test with Dunn’s multiple comparisons test). *p < 0.05, **p < 0.01 and ****p < 0.0001. All data are presented as mean ± SEM.

### Pharmacological suppression of GluN2A- and GluN2B-containing receptors regulate tonic inhibition in the KA-induced seizure model

Finally, we examined whether GluN2-dependent regulation of α5-GABA_A_Rs may occur under pathological conditions. To this end, we induced the acute seizure model by systematic administration of kainic acid (KA), an agonist for kainate- and AMPA-type ionotropic glutamate receptors. All mice injected with KA (20 mg/kg) showed epileptic behavior within 30 min and completely recovered 24 h after injection (Figure S4A). We then analyzed the total and surface expression levels of GluN2A, GluN2B and GABA_A_Rs in hippocampal slices prepared from mice at 1 h or 24 h after KA injection (Figure S4B). We found that KA administration had little effect on the total expression level of NMDAR and GABA_A_R subunits, comparing with saline-injected control (Figures 4A, B and C). In contrast, 1-h post-KA injection produced a significant increase of surface GluN2B, but not GluN2A (Figure 4A and 4B). Interestingly, at 24 h after KA injection, surface GluN2A was increased, whereas surface GluN2B was unaffected (Figure 4A and 4B). As expected from the regulation of surface α5 expression by GluN2A- and GluN2B-containing receptors (Figure 2C), surface expression of α5-GABA_A_R was decreased at 1 h, and then increased at 24 h after KA injection (Figure 4C). In addition, surface α1-GABA_A_R was increased at 24 h after KA injection (Figure S4C). These results suggest the expression of GluN2 subunits and synaptic and extrasynaptic GABA_A_Rs are dynamically regulated by the KA-induced seizure.

**Figure 4.**
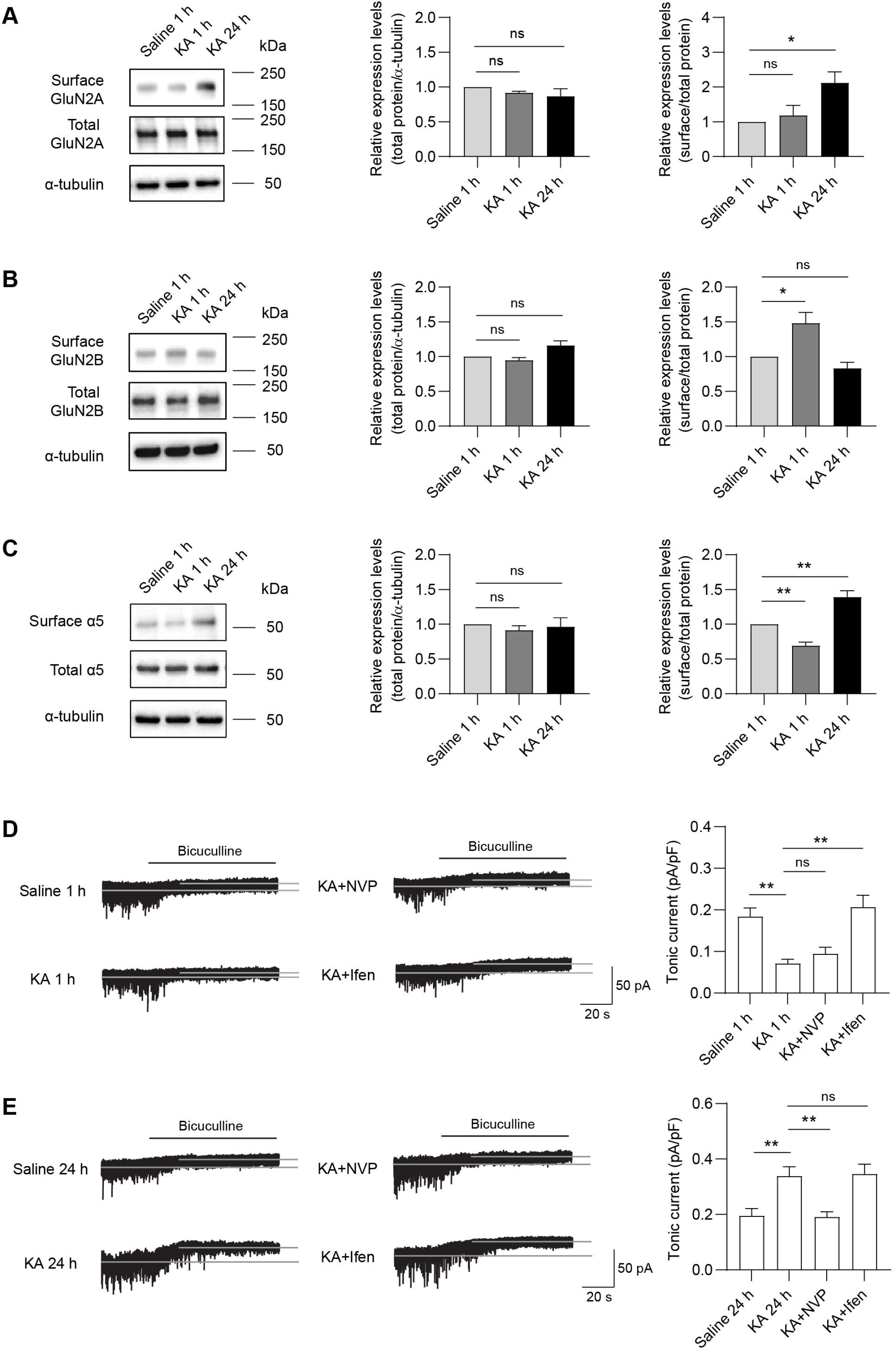
Pharmacological suppression of GluN2A- and GluN2B-containing receptors regulate tonic inhibition in the KA-induced seizure model. (A-C) Representative Western blots and summary graphs from cell-surface biotinylation assays showing that surface and total GluN2A (A), GluN2B (B) and α5-GABA_A_R (C) expression in the KA-induced seizure model. (n = 3 independent experiments, one-way ANOVA with Dunnett’s multiple comparisons test) (D-E) Representative traces (D) and summary graphs (E) showing that tonic currents in hippocampal CA3 neurons were decreased 1 h after KA injection, whereas increased 24 h after KA injection. Ifen or NVP treatment 1 h prior to KA injection respectively restored the decreased or increased tonic currents at corresponding time point after KA injection. (n = 10 for each group, one-way ANOVA with Dunnett’s multiple comparisons test). *p < 0.05 and **p < 0.01. All data are presented as mean ± SEM.

It has been shown that the CA3 region of the hippocampus plays a critical role in KA-induced seizures (Vincent and Mulle, 2009). We thus recorded the tonic currents in CA3 pyramidal cells (Figures S4D). In keeping with the alterations of surface α5 expression after KA injection, we found that whereas tonic currents in hippocampal CA3 neurons were decreased significantly in KA-administrated mice 1 h after induction, they were increased 24 h after induction compared to control mice administrated with saline (Figures 4D and 4E). Interestingly, Ifen (10 mg/kg) or NVP (10 mg/kg) treatment 1 h prior to KA injection respectively restored the decreased or increased tonic currents at corresponding time point after KA injection (Figures 4D and 4E). These results suggest that KA-induced changes of tonic inhibition are dependent on distinct GluN2-containing NMDARs in a temporal-specific manner. In contrast, the mIPSC amplitude was increased 24 h after KA treatment compared with saline treatment, but blockade of GluN2A- or GluN2B-containing NMDARs had little effect on this alteration (Figures S4E and S4F), indicating KA-induced increase of phasic inhibition is independent on GluN2A- and GluN2B-containing NMDARs.

## DISCUSSION

In this study, we investigate the role of GluN2A- and GluN2B-containing NMDARs towards the regulation of tonic inhibition. Our genetic and pharmacological analyses indicate opposing actions of GluN2A- and GluN2B-containing NMDARs on α5-GABA_A_R internalization and tonic inhibition in hippocampal neurons. In addition, GluN2A- but not GluN2B-containing receptors are required for homeostatic potentiation of tonic inhibition. In a KA-induced acute seizure model, while tonic inhibition is decreased during acute seizures, it is increased 24 h later and these alterations are dependent on the activity of the distinct GluN2-containing NMDARs. Collectively, these data extend previous work showing the importance of NMDARs in regulating inhibitory synapse development and transmission (Chiu et al., 2018; Gu and Lu, 2018; Gu et al., 2016; Horn and Nicoll, 2018) and reveal an important crosstalk between glutamatergic signaling and extrasynaptic GABA_A_Rs.

### Differential regulation of tonic inhibition by NMDA receptor subtypes

It has been reported that extrasynaptic GABA_A_R-mediated tonic inhibition is regulated by neurotransmitter receptor-mediated signaling. Indeed, activation of GABA_B_ receptors enhances tonic inhibition in thalamocortical, dentate gyrus, and cerebellar granule cells (Connelly et al., 2013). Additionally, glycine receptors could interact with α5-GABA_A_Rs and regulate tonic currents in the neurons of hypoglossal nucleus (Zou et al., 2019). In addition to inhibitory receptors, tonic inhibition is modulated by glutamatergic receptors expressed at excitatory synapses. For example, genetic deletion of GluN1 leads to augmentation of tonic inhibition in immature hippocampal neurons (Gu et al., 2016). In line with this study, here we found that pharmacological blockade of GluN2B-containing NMDARs, the predominant NMDAR subtype expressed in immature neuron or blockade of all NMDARs enhanced tonic inhibition in immature neurons. Interestingly, we also found that the effect of blockade of GluN2B-containing NMDARs on tonic inhibition disappeared in the more differentiated neurons. The temporal-specific regulation is likely caused by the differential expression of NMDAR GluN2 subunits during development. Indeed, in the hippocampus, during the first 2 weeks after birth, NMDARs undergo a developmental switch from predominantly GluN2B- to GluN2A-containing receptors (Dong et al., 2006). This may also explain the lack of effects on tonic inhibition by blockade of GluN2A-containing receptors in immature neurons. For the 2-3 weeks neurons when both GluN2A and GluN2B are abundantly expressed, opposite actions of GluN2A- and GluN2B-containing receptor blockade on tonic inhibition were observed. Interestingly, inhibition of all NMDARs by APV abolished the differential modulation of tonic inhibition by NMDAR subtypes, presumably due to the normalization of opposing effects of GluN2A- and GluN2B-containing receptors on extrasynaptic GABA_A_Rs. Of note, although both pharmacological inhibition and genetic deletion of GluN2A diminished tonic inhibition in the 2-3 weeks neurons, overexpression of GluN2A had no significant effect on tonic inhibition, showing that further increase in GluN2A expression over endogenous levels at this developmental stage is not sufficient to alter tonic inhibitory currents.

Currently, the molecular mechanisms underlying the regulation of tonic inhibition by NMDARs remain unclear. One scenario is that GluN2A- and GluN2B-containing receptors are coupled to distinct downstream phosphatase and kinase pathways (Shipton and Paulsen, 2014; Sun et al., 2018; Wu and Tymianski, 2018), which in turn may differentially regulate α5-GABA_A_R trafficking and tonic inhibition. For instance, it has been reported that activation of GluN2A- and GluN2B-containing receptors differentially regulates ERK1/2 (Chen et al., 2007) and Akt activities (Liu et al., 2007). In addition, blocking GluN2A-containing receptors attenuates ischemic preconditioning-induced the transcription factor CREB phosphorylation in CA1 pyramidal neurons, whereas blocking GluN2B-containing receptors has little effect (Chen et al., 2008; Terasaki et al., 2010), showing that NMDAR subtypes can activate different signaling pathways. However, it remains unknown whether these molecular pathways are involved in the regulation of α5-GABA_A_R trafficking and tonic inhibition. It has been shown that α5-GABA_A_R trafficking could be regulated in a phosphorylation-dependent manner. Indeed, phosphorylation of radixin, a cytoskeletal protein binding to α5-GABA_A_Rs, increases α5-GABA_A_R clustering at the extrasynaptic sites (Hausrat et al., 2015; Loebrich et al., 2006). In addition, phosphorylation of Shisa7, a GABA_A_R auxiliary subunit (Han et al., 2019), can regulate α5-GABA_A_R trafficking and tonic inhibition (Wu et al., 2021), although Shisa7 primarily modulates α5-GABA_A_R exocytosis. These findings raise the possibility that GluN2A- and GluN2B-containing receptors might differentially regulate tonic inhibition through modulation of phosphorylation of molecules that are involved in α5-GABA_A_R trafficking. It is worth noting that CaMKII activation leads to an increase in cell-surface α5β3-containing receptors and an enhancement of tonic currents (Saliba et al., 2012), and conversely blockade of CaMKII activity causes a reduction of tonic currents in hippocampal neurons (Wu et al., 2021). Interestingly, GluN2A- and GluN2B-containing receptors can differentially regulate CaMKII activity (Strack, McNeill, and Colbran 2000; Barria and Malinow 2005), which in turn might contribute to distinct modulation of α5-GABA_A_R trafficking and tonic inhibition. It has also been reported that activation of GluN2A- and GluN2B-containing NMDARs results in differential expression of brain derived neurotrophic factor (BDNF) triggered by distinct intracellular signaling (Chen et al., 2007). BDNF has been implicated in the regulation of cell-surface expression of extrasynaptic GABA_A_Rs containing α4 and δ subunits in hippocampal neurons (Joshi and Kapur, 2009; Roberts et al., 2006). However, it remains to be determined whether extrasynaptic α5-GABA_A_Rs are similarly modulated by BDNF in an NMDAR subtype-specific manner. In the future, it will be important to determine how GluN2A- and GluN2B-containing NMDARs can distinctly modulate tonic inhibition.

### Alterations in tonic inhibition in epilepsy

It has been reported that GluN2B-containing NMDARs play an important role in the pathophysiology of epilepsy (Waxman and Lynch, 2005; Zhang and Luo, 2013). Specifically, the calcium influx through extrasynaptic GluN2B-containing NMDARs blocks BDNF expression by inactivating CREB and leads to mitochondrial dysfunction and cell death (Hardingham and Bading, 2010; Hardingham et al., 2002). Upregulation of GluN2B mRNA was also found in pyramidal cells of non-sclerotic hippocampi from epileptic patients (Mathern et al., 1998). In an animal model of epilepsy induced by pilocarpin, a cholinergic muscarinic agonist, surface accumulation of GluN2B-containing NMDARs in the hippocampus has been reported (Muller et al., 2013; Naylor et al., 2013). In addition, it has been shown that a decrease in α5-containing GABA_A_Rs occurs in CA1 pyramidal cells of pilocarpine-treated rats (Houser and Esclapez, 2003). In this study, we show that KA injection induces upregulation of surface GluN2B and downregulation of surface α5-GABA_A_R in the hippocampus 1 h after KA injection. Together with the experiments demonstrating that GluN2B-containing NMDARs regulate α5-GABA_A_R internalization, our data indicate that an increase of GluN2B-containing receptors contributes to the reduction of surface α5-GABA_A_R by enhancing endocytosis, resulting in downregulation of tonic inhibition during acute seizures. Given that the important role of tonic inhibition in regulating neural network excitability (Belelli et al., 2009; Farrant and Nusser, 2005; Lee and Maguire, 2014), GluN2B receptor-mediated reduction of tonic inhibition may contribute to KA-induced neuronal hyperexcitability and epileptogenesis in the seizure model.

Our data also show that in the KA-induced acute seizure model, whereas epileptic phenotypes disappear 24 h after KA induction, tonic inhibition in hippocampal CA3 neurons is increased. Consistently, a recent study using a similar seizure model has shown that enhanced neuronal excitability in cortical neurons returns to the basal level and tonic inhibition is increased 24 h after KA injection (Pan et al., 2018). Additionally, in the pilocarpine-induced seizure model, the tonic current is enhanced in dentate gyrus granule cells 3 weeks after induction (Naylor et al., 2005). Thus, an increase of tonic inhibition after acute seizure appears to be a common phenotype across different neuronal types in different seizure models. What is the functional significance of the increased tonic current 24 h later after acute seizures? Considering the importance of tonic GABAergic signaling in regulating neural network excitability (Belelli et al., 2009; Farrant and Nusser, 2005; Lee and Maguire, 2014), the increase in tonic inhibition that occurs after acute seizures may act as a homeostatic adaptive response to KA-induced neuronal hyperexcitability, and thus may help lessen neurodegeneration associated with neuronal hyperexcitability. We also found that homeostatic potentiation is dependent on the GluN2A subunit. In the adult forebrain, GluN2A-containing receptors are predominantly localized at synaptic sites and activation of synaptic NMDARs is considered to be neuroprotective (Hardingham and Bading, 2010; Hardingham et al., 2002). Therefore, GluN2A-dependent upregulation of tonic inhibition after KA-induced seizure may provide additional evidence supporting the neuroprotective effects of GluN2A-containing receptors in neurodegenerative disorders (Liu et al., 2007; Terasaki et al., 2010). In addition, in line with a previous report (Peng et al., 2010), we found that the phasic inhibition could undergo the homeostatic potentiation after KA administration.

However, homeostatic potentiation of synaptic inhibition is independent of GluN2A, suggesting distinct mechanisms governing homeostatic plasticity of tonic and phasic inhibition induced by KA.

Taken together, we have uncovered a critical role of NMDARs in the regulation of α5-GABA_A_R trafficking and tonic inhibition during development and in a seizure model, and revealed distinct roles of NMDAR subtypes in regulating tonic inhibition in a development-specific manner. As dysregulations of tonic inhibition has been shown to be a mechanism underlying a variety of pathological brain states (Brickley and Mody, 2012; Hines et al., 2012), our findings also offer insight into developing more selective treatment of a diverse array of neurological and psychiatric disorders.

## Acknowledgements

We are grateful to all members from Wei Lu laboratory for critical comments on the manuscript. This work was supported by the NIH/NINDS Intramural Research Program (W.L.).

## Author Contributions

K.W. and W.L. designed the project, and W.L. supervised the project. K.W. performed imaging, biochemical and behavioral experiments, and electrophysiological recordings. K.W. and D.C. cloned sgRNA constructs. Q.T. performed neuronal cultures. W.L. and K.W. wrote the manuscript, and all authors read and commented on the manuscript.

## STAR METHODS

### Resource Availability

#### Lead contact

Further information and requests for resources and reagents should be directed to the Lead Contact, Wei Lu.

### Materials Availability

All unique reagents generated in this study are available from the Lead Contact with a completed Materials Transfer Agreement.

### Data and Code Availability

This study did not generate datasets/code.

## EXPERIMENTAL MODEL AND SUBJECT DETAILS

### Animals

All animal handling was performed in accordance with animal protocols approved by the Institutional Animal Care and Use Committee (IACUC) at NIH/NINDS. All mice were housed and bred in a conventional vivarium with ad libitum access to food and water under a 12-h circadian cycle. Time-pregnant mice at E17.5-18.5 were used for dissociated hippocampal neuronal culture. Young adult male mice (6-8 weeks old) were used for biochemical, electrophysiological, and behavioral experiments.

### Cell Lines

HEK293T cells (ATCC, Cat# CRL-11268) were maintained with culture media containing 1% penicillin-streptomycin (GIBCO), 10% FBS (GIBCO) in Dulbecco’s Modified Eagle’s Medium (DMEM, GIBCO), in a humidified incubator at 37 °C with 5% CO_2_.

### Dissociated Hippocampal Neuronal Culture

Mice hippocampal neurons were prepared from E17.5-18.5 mice embryos of either sex as previously described (Wu et al., 2021). In brief, the hippocampi were dissected from embryonic brains and digested in the Hank’s Balanced Salt Solution (HBSS, GIBCO) containing 20 U/ml papain (Worthington) and 100 U/ml DNase I (Worthington) at 37 °C for 45 min. After centrifugation for 5 min at 800 rpm, the pellet was resuspended in HBSS containing 100 U/ml DNase I, and was fully dissociated by pipetting up and down. Cells were then transferred into HBSS containing trypsin inhibitor (10 mg/ml, Sigma-Aldrich) and BSA (10 mg/ml, Sigma-Aldrich). After centrifugation for 10 min at 800 rpm, cells were resuspended in Neurobasal media (GIBCO) supplemented with 2% B27 (GIBCO) and 2 mM GlutaMAX (GIBCO) and were plated on poly-D-lysine (Sigma-Aldrich)-coated glass coverslips or 6-well plates. Cultures were maintained in Neurobasal media supplemented with 2% B27 and 2 mM GlutaMAX in a humidified incubator at 37 °C with 5% CO_2_. Culture media were changed by half volume once a week.

## METHOD DETAILS

### Plasmids

pRK5-GFP-GluN2A and pRK5-GFP-GluN2B were gifts from Katherine Roche’s lab at NINDS, NIH. pSpCas9(BB)-2A-Puro (PX459) V2.0 and pSpCas9(BB)-2A-GFP (PX458) were purchased from Addgene. Custom oligonucleotides were generated (GluN2A forward, 5’ CACCGCGACGTGACAGAACGCGAAC 3’; and GluN2A reverse, 5’ AAACGTTCGCGTTCTGTCACGTCGC 3’; and GluN2B forward, 5’ CACCGTCTGACCGGAAGATCCAGG 3’; and GluN2B reverse, 5’ AAACCCTGGATCTTCCGGTCAGAC 3’; IDT), and cloned into pSpCas9-BB-2A-GFP (PX458) or pSpCas9(BB)-2A-Puro (PX459) V2.0 vector at the BbsI cutting site. The coding sequence of GluN2A and GluN2B point mutations for sgRNA resistant plasmid (GluN2A: AGCCACGACGTGACAGAACGCGAACTT to AG<u>T</u>CACGACGTGAC<u>T</u>GA<u>GA</u>G<u>A</u>GAACTT; GluN2B: ATGTCTGACCGGAAGATCCAGGGG to ATGTCTGA<u>T</u>CG<u>T</u>AAGAT<u>T</u>CA<u>A</u>GG<u>A</u>) were generated by Q5 Site-Directed Mutagenesis Kit (NEB).

### Cell Transfection

HEK-293T cells were transfected with GluN2A or GluN2B, together with sgRNA using CalPhos Mammalian Transfection Kit (Takara). Western blot was performed 48 h after transfection. Hippocampal neurons at DIV3-4 were transfected with GluN2A sgRNA or GluN2B sgRNA using NeuroMag reagent (Oz Biosciences), and were recorded at DIV16-17. Hippocampal neurons at DIV11 were co-transfected with pCAG-IRES-GFP and GluN2A or GluN2B using NeuroMag reagent. Electrophysiological recordings or immunostaining were performed 72 h after transfection. All transfection kits were used according to the manufacturer’s instructions.

### Electrophysiology

For recording in dissociated hippocampal cultures, neurons were continuously perfused with the extracellular solution containing (in mM): 140 NaCl, 5 KCl, 2 CaCl_2_, 1 MgCl_2_, 10 HEPES, and 10 Glucose (pH 7.3; osmolality 300-310 mOsm). The internal solution contained (in mM): 70 CsMeSO4, 70 CsCl, 8 NaCl, 10 HEPES, 0.3 Na-GTP, 4 Mg-ATP and 0.3 EGTA (pH 7.3; osmolality 285-290 mOsm). Miniature inhibitory postsynaptic currents (mIPSCs) and tonic currents were recorded at −70 mV in the presence of 0.5 μM TTX (Alomone Labs) and 20 μM DNQX (Alomone labs). For recording NMDA mEPSCs at +40 mV, 0.5 μM TTX, 20 μM DNQX, and 50 μM picrotoxin (Sigma-Aldrich) were added into the extracellular solution. For recording NMDA-induced whole-cell currents, TTX (0.5 μM) were added into 0 Mg^2+^ extracellular solution. NMDA-induced current was recorded at −70 mV by rapid application/removal of NMDA (100 μM) using a computer-controlled multi-barrel perfusion system (Automate Scientific).

For recording in acute brain slices, transverse hippocampal slices (300 μm thickness) were prepared from 6-8 weeks old male mice in chilled high sucrose cutting solution, containing (in mM): 2.5 KCl, 0.5 CaCl_2_, 7 MgCl_2_, 1.25 NaH_2_PO_4_, 25 NaHCO_3_, 7 glucose, 210 sucrose and 1.3 ascorbic acid. The slices were recovered in artificial cerebrospinal fluid (ACSF) containing (in mM): 130 NaCl, 3.5 KCl, 24 NaHCO_3_, 1.25 NaH_2_PO_4_-H_2_O, 10 glucose, 2.5 CaCl_2_ and 1.5 MgCl_2_ (pH 7.3; osmolality 300-310 mOsm) at 33 °C for 30 min and then were maintained at room temperature prior to recording. To record tonic currents, slices were transferred to a submersion chamber, continuously perfused with ACSF with 0.5 μM TTX, 20 μM DNQX and 5 μM GABA. The intracellular solution contained (in mM) 130 CsCl, 8.5 NaCl, 5 HEPES, 4 MgCl_2_, 4 Na-ATP, 0.3 Na-GTP and 1 QX-314 (pH 7.3; osmolality 285-290 mOsm).

To measure tonic inhibitory currents in neuronal cultures or in acute hippocampal slices, the GABA_A_R competitive antagonist bicuculline (20 μM, Abcam) was bath applied after obtaining a stable baseline recording at −70 mV. An all-points histogram was plotted for a 20-s period before and during bath-application of bicuculline, fitting the histogram with a Gaussian distribution gave the mean baseline holding currents, and the difference in baseline holding currents before and during bicuculline application was calculated to be the tonic currents. Tonic currents were normalized to membrane capacitance, to account for variability in cell size. Series resistance was monitored and not compensated, and cells in which series resistance was more than 25 MΩ or varied by 25% during a recording session were discarded. Whole-cell recordings were obtained from cells visualized with a fixed stage upright microscope (BX51WI, Olympus). Fluorescence-positive cells were identified by epifluorescence microscopy. Data were collected with a Multiclamp 700B amplifier (Axon Instruments), filtered at 2 kHz, and digitized at 10 kHz.

### Immunostaining

For surface α5 receptor labeling, cultured hippocampal neurons at DIV15 on coverslips were incubated with anti-α5 antibody (1:500, Synaptic Systems) in culture medium for 15 min. Next, they were washed briefly with fresh culture medium and fixed with a solution containing 4% paraformaldehyde and 4% sucrose in PBS. Cultured neurons were subsequently incubated with Alexa 555-conjugated anti-rabbit secondary antibody (1:1000, Thermo Fisher Scientific) for the visualization of α5. For the endocytosis assay, cultured hippocampal neurons at DIV15 were incubated live with rabbit α5 antibody (1:500, Synaptic Systems) at 37°C for 10 min in conditioned culture medium. After incubation, the neurons were washed with PBS and then incubated in antibody-free medium to allow antibody-bound receptors to undergo internalization at 37°C for 30 min, followed by fixation with 4% paraformaldehyde and 4% sucrose in PBS. After fixation, neurons were washed and then blocked with 10% NGS for 1 h, exposed to Alexa 488-conjugated anti-rabbit secondary antibody (1:200, Jackson ImmunoResearch Labs) for 1 h under the nonpermeabilized condition, and then internalized α5 was labelled with Alexa 555-conjugated anti-rabbit secondary antibody (1:1000, Thermo Fisher Scientific) for 1h after permeabilization in PBS containing 0.25% Triton X-100 and blocking in 10% NGS. Coverslips were washed for three times with PBS and mounted with Fluoromount-G.

Fluorescence images were acquired on a Zeiss LSM 880 laser scanning confocal microscope with a 63 x 1.4 NA oil immersion objective. For quantification, sets of cells were prepared and stained simultaneously. Compared images were acquired at the same time using identical acquisition settings. The fluorescence intensity was analyzed using ImageJ.

### Surface cell biotinylation of hippocampal slices

Hippocampal slices were prepared from 6-8 weeks mice as described (Li et al., 2017). Surface expression of GluN2A, GluN2B, α1-GABA_A_R and α5-GABA_A_R was quantitated as described. Briefly, acute hippocampal slices were labeled for 30 min at 4 °C with 1 mg/ml sulfo-NHS-SS biotin (ThermoFisher Scientific). Membranes were prepared and the biotinylated proteins were precipitated with streptavidin agarose resin (ThermoFisher Scientific) and detected by western blot.

### Kainic acid-induced seizure model

Young adult male mice (6-8 weeks old) were administered an intraperitoneal (i.p.) injection of kainic acid (KA, Abcam) dissolved in 0.9% saline solution at 20 mg/kg body weight. Seizure score was evaluated at 0 h, 0.5 h, 1 h and 24 h after KA injection according to the modified Racine scale (Racine, 1972): stage 0, normal behavior; stage 1, immobility and rigidity; stage 2, repetitive behaviors, head nodding or bobbing; stage 3, Forelimb clonus with partial or intermittent rearing; stage 4, continuous rearing and falling; stage 5, severe clonic-tonic seizures; stage 6, death. The expression levels of GluN2A, GluN2B and GABA_A_Rs in hippocampi were examined at 1 h or 24 h after KA injection. To examine the effects of GluN2A- and GluN2B-containing receptors on the tonic inhibition in KA-induced seizure model, mice were injected with NVP (10 mg/kg), Ifen (10 mg/kg) or saline 1 h prior to KA injection and were sacrificed for electrophysiological recordings at 1 h or 24 h after KA injection.

## QUANTIFICATION AND STATISTICAL ANALYSIS

### Statistical Analysis

Statistical analysis was performed in GraphPad Prism 8.0 software. Normality distribution was tested by the Shapiro-Wilk test before carrying out a subsequent statistical test. Direct comparisons between two groups were made using two-tailed Student’s t test or Mann-Whitney U test. Multiple comparisons were performed using one-way ANOVA, Kruskal-Wallis test or two-way ANOVA with corrections for multiple comparisons test (see figure legends for specifics).

