## Supplementary figures for "Distinct regulation of tonic GABAergic inhibition by NMDA receptor subtypes"

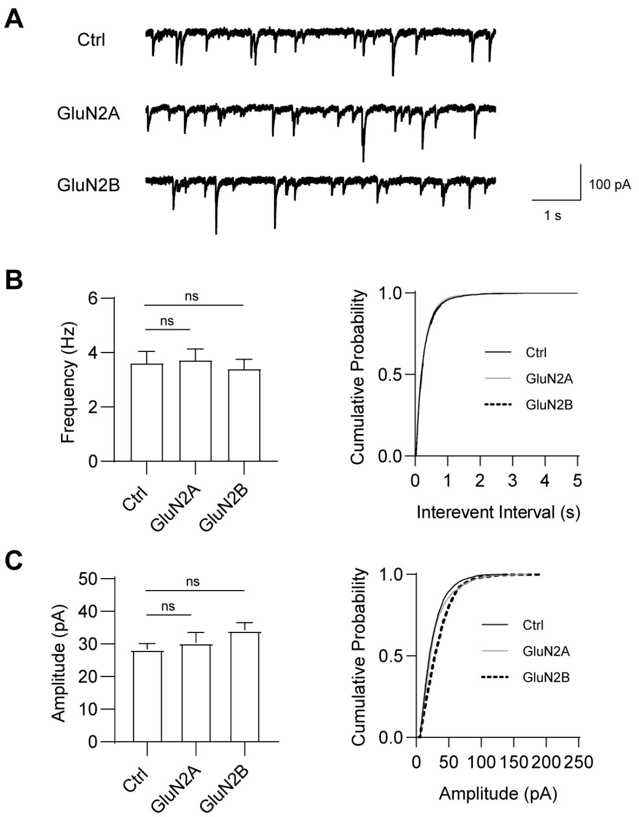


**FigS1. Overexpression of GluN2A or GluN2B has no effect on phasic inhibition**

(A) Representative mIPSCs traces recorded in cultured hippocampal neurons expressing pCAG-GFP alone or pCAG-GFP together with GluN2A or GluN2B.

(B) mIPSCs mean frequency and cumulative probability plots of mIPSC interevent intervals. (n = 11-14 for each group, one-way ANOVA with Dunnett's multiple comparisons test).

(C) mIPSCs mean amplitude and cumulative probability plots of mIPSC amplitude. (n = 11-14 for each group, one-way ANOVA with Dunnett's multiple comparisons test).

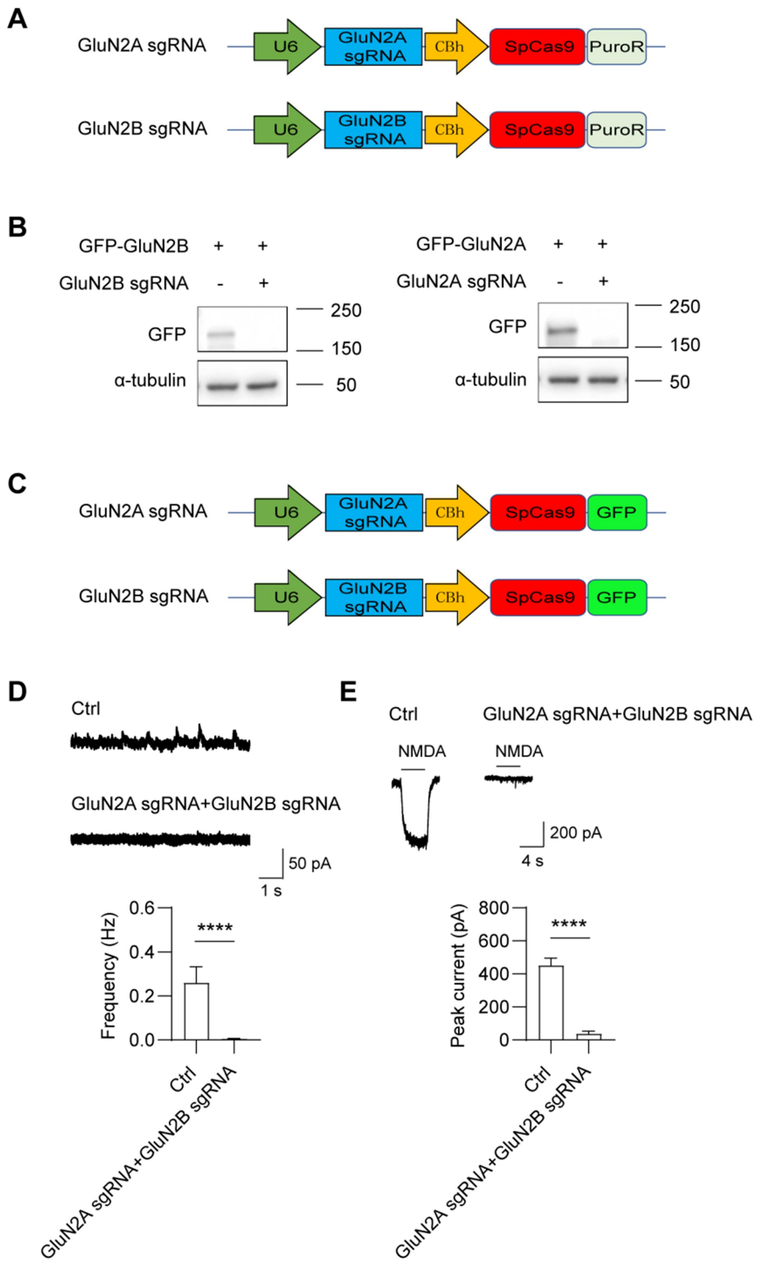
A
GluN2A sgRNA
U6
GluN2A sgRNA
CBh
SpCas9
PuroR
GluN2B sgRNA
U6
GluN2B sgRNA
CBh
SpCas9
PuroR
B
GFP-GluN2B + +
GluN2B sgRNA - +
GFP
α-tubulin
250
150
50
GFP-GluN2A + +
GluN2A sgRNA - +
GFP
α-tubulin
250
150
50
C
GluN2A sgRNA
U6
GluN2A sgRNA
CBh
SpCas9
GFP
GluN2B sgRNA
U6
GluN2B sgRNA
CBh
SpCas9
GFP
D
Ctrl
GluN2A sgRNA+GluN2B sgRNA
50 pA
1 s
Frequency (Hz)
0.6
0.4
0.2
0.0
****
Ctrl
GluN2A sgRNA+GluN2B sgRNA
E
Ctrl
GluN2A sgRNA+GluN2B sgRNA
NMDA
NMDA
200 pA
4 s
Peak current (pA)
800
600
400
200
0
****
Ctrl
GluN2A sgRNA+GluN2B sgRNA

**FigS2. Validation of knockout efficacy of GluN2A and GluN2B sgRNAs**

(A) Schematic diagram of CRISPR/Cas9 vector (without GFP tag) targeting GluN2A or GluN2B gene used for biochemical experiments.

(B) Representative Western blots of HEK293 cell lysates showing expression of GluN2A and GluN2B following co-transfection of either GFP-GluN2A or GFP-GluN2B together with the corresponding sgRNA as shown in A.

(C) Schematic diagram of CRISPR/Cas9 vector (with GFP tag) targeting GluN2A or GluN2B gene used for electrophysiological experiments.

(D) NMDA mEPSCs recorded in cultured hippocampal neurons transfected with GluN2A gRNA and GluN2B sgRNA as shown in C. (n = 9-10 for each group, Mann-Whitney U test).

(E) NMDA-evoked currents recorded in cultured hippocampal neurons transfected with GluN2A sgRNA and GluN2B sgRNA. (n = 9-16 for each group, Mann-Whitney U test).

*$p < 0.05$ and **$p < 0.01$. All data are presented as mean ± SEM.

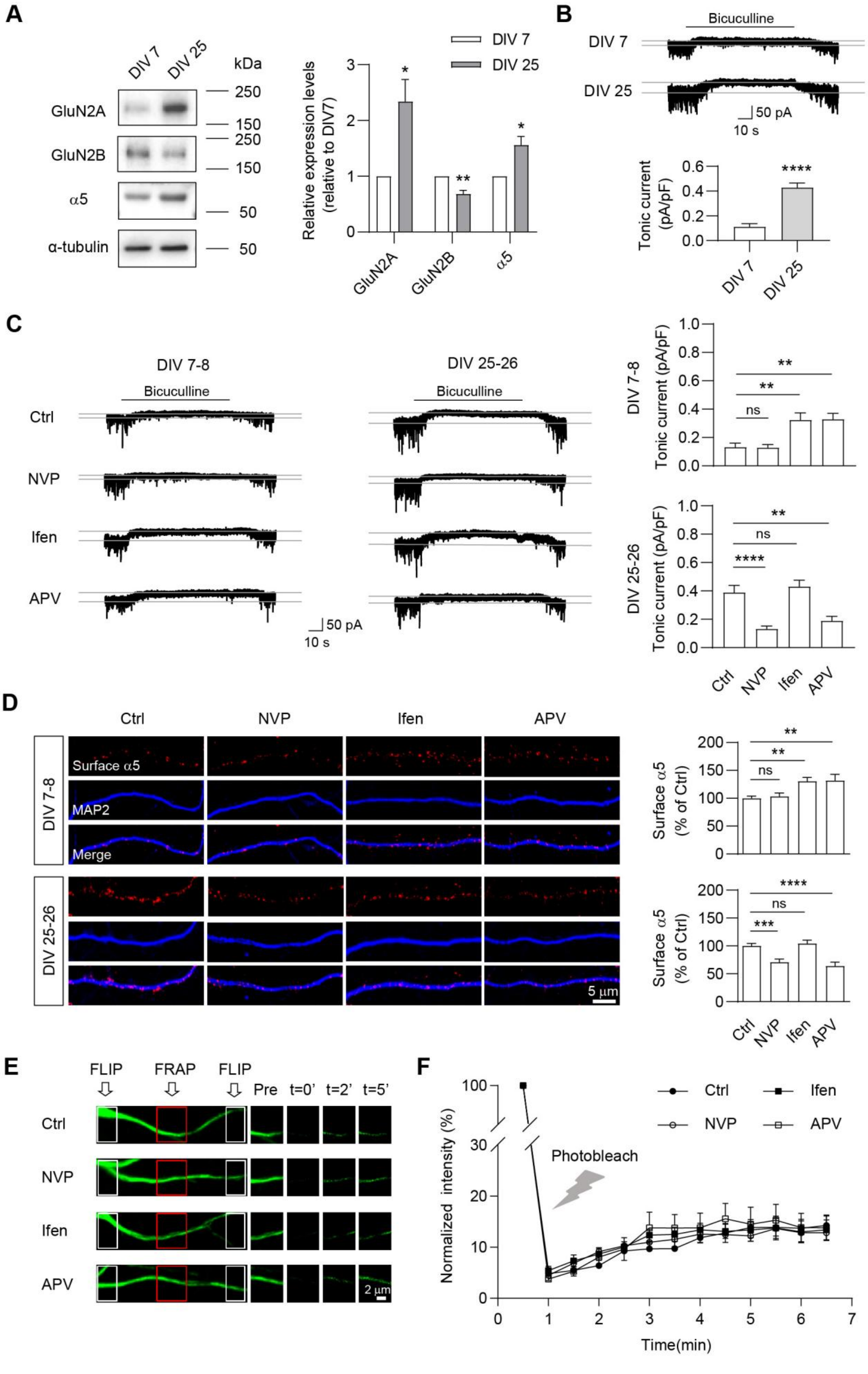
A
DIV 7
DIV 25
kDa
GluN2A
GluN2B
α5
α-tubulin
250
150
250
150
50
50
Relative expression levels (relative to DIV7)
GluN2A
GluN2B
α5
B
Bicuculline
DIV 7
DIV 25
50 pA
10 s
Tonic current (pA/pF)
****
C
DIV 7-8
DIV 25-26
Bicuculline
Ctrl
NVP
Ifen
APV
50 pA
10 s
DIV 7-8 Tonic current (pA/pF)
DIV 25-26 Tonic current (pA/pF)
ns
**
****
D
Surface α5
MAP2
Merge
5 μm
Surface α5 (% of Ctrl)
***
E
FLIP
FRAP
Pre
t=0'
t=2'
t=5'
2 μm
F
Normalized intensity (%)
Photobleach
Time(min)

**FigS3. Development-dependent regulation of tonic inhibition by GluN2A- and GluN2B-containing receptors**

(A) Representative Western blots and summary graphs showing the expression level of GluN2A, GluN2B and $\alpha$5-GABA$_A$R in the homogenates of hippocampal neurons at DIV7 and DIV25. (n = 3 independent experiments, t test)

(B) Representative traces and summary graphs showing tonic currents in cultured hippocampal neurons at DIV7 and DIV25. (n = 8-10 for each group, Mann-Whitney U test)

(C) Representative traces and summary graphs showing the changes of tonic currents in immature (DIV7-8) or more differentiated (DIV25-26) hippocampal neurons under treatments of NVP, Ifen or APV. (n = 10-11 for each group, one-way ANOVA with Dunnett's multiple comparisons test)

(D) Immunostaining and summary graphs showing changes of surface $\alpha$5 expression in immature or more differentiated hippocampal neurons under treatments of NVP, Ifen or APV. (n = 22-39 for each group, Kruskal-Wallis test with Dunnett's multiple comparisons test)

(E) Representative images (left) of SEP-$\alpha$5 fluorescence and the regions of the neuronal dendrites used for the fluorescence recovery after photobleaching (FRAP) and fluorescence loss in photobleaching (FLIP) experiments. Repetitive photobleaching (FLIP, white box) occurred at regions bilateral to the central FRAP region (red box). Each column (right) represents before (pre), immediately after (t = 0'), and at 2 min (t =2') and 5 min (t = 5') after photobleaching in each condition.

(F) Normalized fluorescence recovery curves showing that NVP, Ifen and APV had no effects on $\alpha$5 exocytosis. (n = 6 for each group, two-way ANOVA with Dunnett's multiple comparisons test)

*$p < 0.05$, **$p < 0.01$, ***$p < 0.001$ and ****$p < 0.0001$. All data are presented as mean ± SEM.

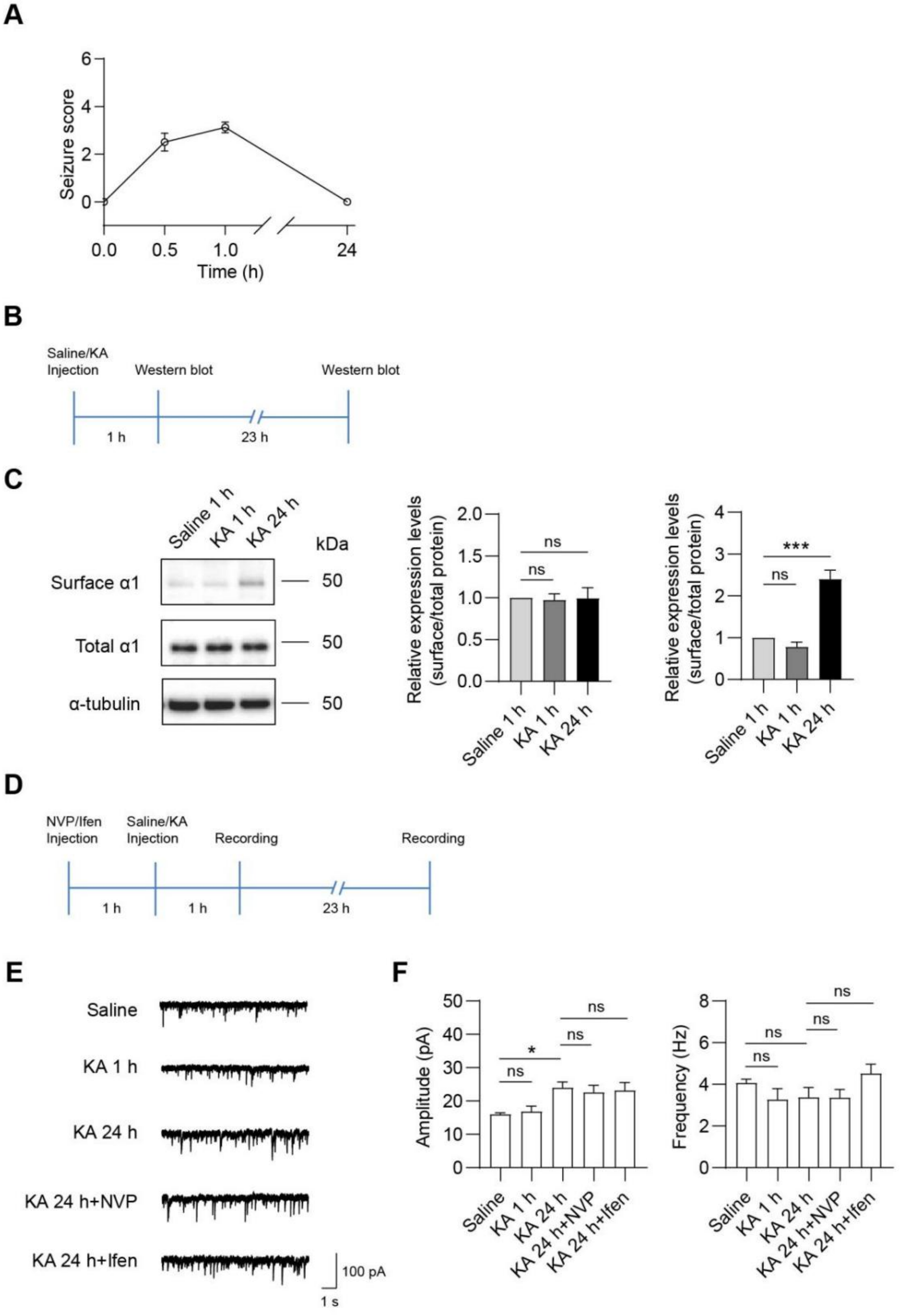
A
Seizure score
Time (h)
0.0
0.5
1.0
24
B
Saline/KA Injection
Western blot
Western blot
1 h
23 h
C
Saline 1 h
KA 1 h
KA 24 h
kDa
Surface α1
Total α1
α-tubulin
50
Relative expression levels (surface/total protein)
ns
***
D
NVP/Ifen Injection
Saline/KA Injection
Recording
Recording
1 h
1 h
23 h
E
Saline
KA 1 h
KA 24 h
KA 24 h+NVP
KA 24 h+Ifen
100 pA
1 s
F
Amplitude (pA)
Frequency (Hz)
ns
*
Saline
KA 1 h
KA 24 h
KA 24 h+NVP
KA 24 h+Ifen

**FigS4. Pharmacological suppression of GluN2A- and GluN2B-containing receptors have no effect on phasic inhibition in hippocampal CA3 neurons in a KA-induced seizure model**

(A) Seizure score was evaluated at 0 h, 0.5 h, 1 h and 24 h after KA injection according to the modified Racine scale.(n = 8)

(B) Experimental design for Western blot.

(C) Representative Western blots and summary graphs from cell-surface biotinylation assays showing surface and total $\alpha$1-$GABA_AR$ expression in KA-induced seizure model. (n = 3 independent experiments, one-way ANOVA with Dunnett's multiple comparisons test)

(D) Experimental design for electrophysiological recording.

(E) Representative mIPSCs traces recorded in hippocampal CA3 neurons in acute brain slices.

(F) Summary graphs showing mean amplitude and frequency of mIPSCs. (n = 10-16 for each group, one-way ANOVA with Tukey's multiple comparisons test).

*$p < 0.05$ and ***$p < 0.001$. All data are presented as mean ± SEM.
